# LiPyphilic: A Python toolkit for the analysis of lipid membrane simulations

**DOI:** 10.1101/2021.05.04.442445

**Authors:** Paul Smith, Christian D. Lorenz

**Affiliations:** Department of Physics, King’s College London, London, WC2R 2LS, UK

## Abstract

Molecular dynamics simulations are now widely used to study emergent phenomena in lipid membranes with complex compositions. Here, we present LiPyphilic - a fast, fully tested, and easy to install Python package for analysing such simulations. Analysis tools in LiPyphilic include the identification of cholesterol flip-flop events, the classification of local lipid environments, and the degree of interleaflet registration. LiPyphilic is both force field and resolution agnostic, and thanks to the powerful atom selection language of MDAnalysis it can handle membranes with highly complex compositions. LiPyphilic also offers two on-the-fly trajectory transformations to i) fix membranes split across periodic boundaries and ii) perform nojump coordinate unwrapping. Our implementation of nojump unwrapping accounts for fluctuations in box volume under the NPT ensemble — an issue that most current implementations have overlooked. The full documentation of LiPyphilic, including installation instructions, is available at https://lipyphilic.readthedocs.io/en/latest.

**Graphical TOC Entry:** 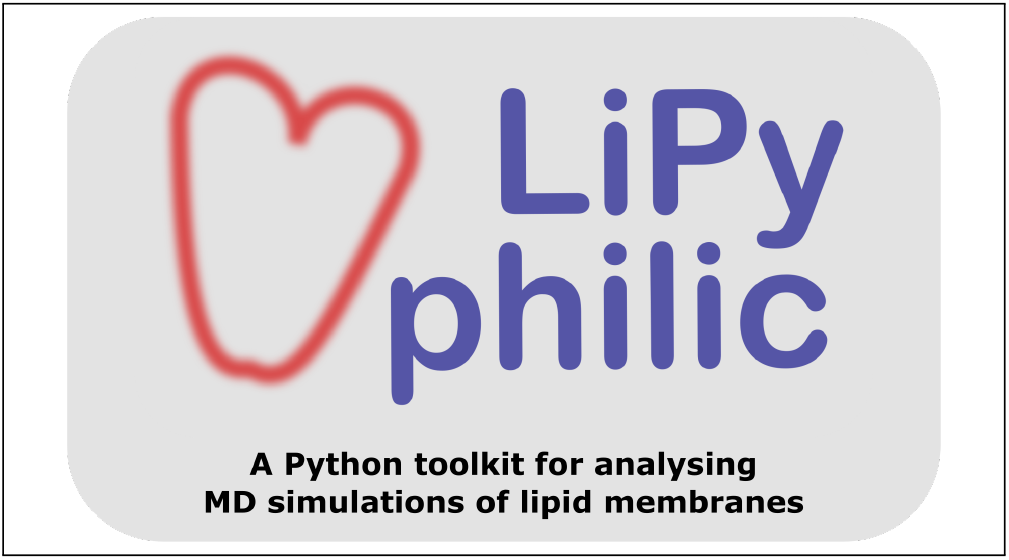

## 1 Introduction

The plasma membrane was once thought to be a passive divider between a cell and its external environment. We now understand that it is in fact a dynamic interface upon which many cellular processes, from cell-signaling to membrane transport, depend. These processes are emergent phenomena that arise from a complex interplay between the molecular species that comprise the plasma membrane. As such, there is a great interest in understanding how the lipids, proteins, and carbohydrates of the plasma membrane interact with one another. Molecular dynamics (MD) simulations are routinely used to study lipid-lipid and lipid-protein interactions at a molecular level, and there exists many excellent tools for analysing the trajectories of such simulations. FATSLiM^1^ and MemSurfer,^2^ for example, both specialise in the analysis of non-planar membranes such as buckled bilayers or vesicles. PyLipID^3^ and ProLint^4^ are designed for the easy and efficient analysis of lipid-protein interactions. ML-LPA is a recently developed Python package that employs various machine learning algorithms to identify the phase — *L*_*o*_ or *L*_*d*_ — of lipids in a bilayer.^5^ LOOS, on the other hand, is a C++ library with a Python interface for analysing MD simulations.^6,7^ Unlike the above packages, LOOS handles the trajectory reading internally whilst also offering a large set of analysis tools, some of which are for lipid membranes. Between them, these software packages provide an extensive analysis suite for MD simulations of lipid membranes.

There are, however, some non-trivial analyses that are frequently employed but are not yet available in any analysis software we are aware of. These include the identification of cholesterol flip-flop events,^8–26^ the classification of local lipid environments,^17,27–34^ and calculating the degree of interleaflet registration. ^11,34–42^ These analyses provide important information about the structure and dynamics of lipid membranes, but they currently require the writing of in-house scripts. Here, we present LiPyphilic — a fast, fully tested, and easy to install Python package that can perform these analyses, among others.

## 2 LiPyphilic

LiPyphilic is an object-oriented Python package for analysing MD simulations of lipid membranes. It is built directly on top of MDAnalysis, and makes use of NumPy^43^ and SciPy^44^ for efficient computation. It is both resolution and force field agnostic, working with any file format that MDAnalysis can load so long as the topology contains residue names. All analysis classes in LiPyphilic share the same application programming interface (API) as those in MDAnalysis. This shared API makes it simple for users of MDAnalysis to learn how to use LiPyphilic.

At its core, LiPyphilic is designed to easily integrate with the wider scientific Python stack. Results are typically stored in a two dimensional NumPy array of shape *N*_*lipids*_ by *N*_*frames*_ (Figure 1), making it simple to post-process the results for further analysis. Some analysis tools also take a two dimensional NumPy array of the same shape as input. This input array may contain information about each lipid, such as which leaflet they belong to or their phase (*L*_*o*_ or *L*_*d*_). Alternatively, the input array may be a boolean mask — an array of True and False values specifying which lipids to include in the analysis. As these inputs are generic NumPy arrays instead of types specific to LiPyphilic, it is possible to use the output from other membrane analysis tools as input to LiPyphilic. For example, you may assign lipids to leaflets using FATSLiM,^1^ determine their phase state using ML-LPA, ^5^ or calculate local membrane normals using MemSurfer, ^2^ and extract the results to perform further analysis with LiPyphilic.

**Figure 1:**
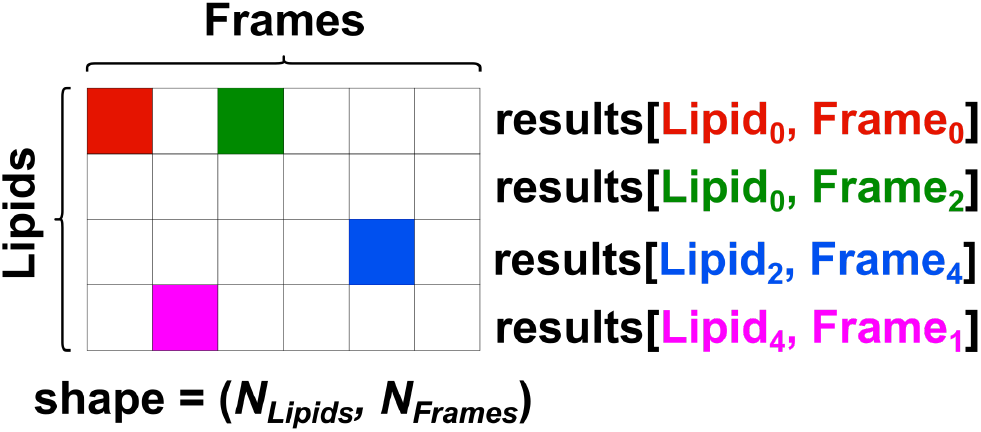
In LiPyphilic, analysis results are typically stored in a NumPy array of shape (*N*_*Lipids*_, *N*_*frames*_)

The workflow for using LiPyphilic generally involves the following steps:

1. import MDAnalysis along with the required LiPyphilic analysis modules
2. load a topology and trajectory as an MDAnalysis universe
3. create an analysis object using the MDAnalysis universe and specifying the relevant input options
4. use the `run()` method to perform the analysis
5. store the results, either by serialising the analysis object itself or saving the results data as a NumPy array

Below we will discuss the implementation and usage of some of the analysis tools currently available in LiPyphilic. We will then discuss the on-the-fly transformations that LiPyphilic can perform on MDAnalysis trajectories. We then provide benchmarks of the analysis tools and transformations. Finally, we will briefly discuss the software engineering best practices used in developing LiPyphilic. If you are more interested in learning how to use LiPyphilic, rather than how LiPyphilic works per se, we recommend working through the examples in the documentation at https://lipyphilic.readthedocs.io/en/latest/reference/analyses.html.

### 2.1 Assign leaflets

For many analyses, such as calculating the area per lipid, it is necessary to know the leaflet within which a lipid is found. LiPyphilic has two tools for assigning lipids to lealfets. The class `lipyphilic.lib.assign_leaflets.AssignLeaflets` assigns each lipid to a leaflet based on the distance in *z* to its local membrane midpoint. This is suitable only for planar bilayers. On the other hand, the class `lipyphilic.lib.assign_leaflets.AssignCurvedLeaflets` can be used to identify leaflets in a buckled bilayer or a micelle. This uses the MDAnalysis leaflet finder ^45,46^ to assign non-translocating lipids to leaflets, then at each frame assigns the remaining lipids based on their minimum distance to each leaflet. `AssignLeaflets` remains useful for planar bilayers, especially if the rate of cholesterol translocation is of interest, which is typically measured by assigning lipids to leaflets based on their *z*-coordinate.

LiPyphilic can assign molecules not just to the upper or lower leaflet, but also to the midplane. This is useful for studying, for example, the local lipid environment of midplane cholesterol^24^ or its role in the registration of nanodomains. ^47^ Assigning cholesterol to the midplane also creates a buffer zone for determining whether a flip-flop event was successful (i.e. crossed the buffer zone) or not.^25^

`AssignLeaflets` and `AssignCurvedLeaflets` have opposing approaches to assigning molecules to the midplane. The former considers a molecule’s distance to the midplane whereas the latter considers its distance to each leaflet. There is naturally an inverse relationship between these two measures - the further a molecule is from the midplane the closer it is to a leaflet. However, the distance to the midplane is typically employed when studying flip-flop in planar bilayers,^17,21–25,31^ whereas the distance to each leaflet is used for studying flip-flop in undulating bilayers.^47^ As with all analysis tools in LiPyphilic, the assigning of lipids to leaflets is both resolution and force field agnostic. Instead of reading atom selections hard coded into the package, the analysis tools rely on the powerful selection language of MDAnalysis. Figure 2A shows how `AssignLeaflets` may be used to determine the leaflet membership of all lipids in the 58 component neuronal plasma membrane studied by Ingólfsson *et alii*.^31^ First, an MDAnalysis Universe must be created. Lipids are then assigned to leaflets by passing this Universe to `AssignLeaflets` along with an atom selection of lipids in the bilayer, using the `universe` and `lipid_sel` arguments respectively. Optionally, to allow molecules to be in the midplane, we can use the `midplane_sel` and `midplane_cutoff` arguments. In the example shown in Figure 2A, cholesterol will be assigned to the midplane if its ROH bead is within 8 Å of its local midpoints. Local midpoints are computed by first splitting the membrane into an *n* by *n* grid in *xy*, where *n* is specified using the `n_bins` argument. The local midpoint of a grid cell is then given by the center of mass of all atoms selected by `lipid_sel` that are in the grid cell. Through calculating local membrane midpoints, this algorithm can account for small undulations in a bilayer. However, for bilayers with large undulations or for non-bilayer membranes, `AssignCurvedLeaflets` should be used.

**Figure 2:**
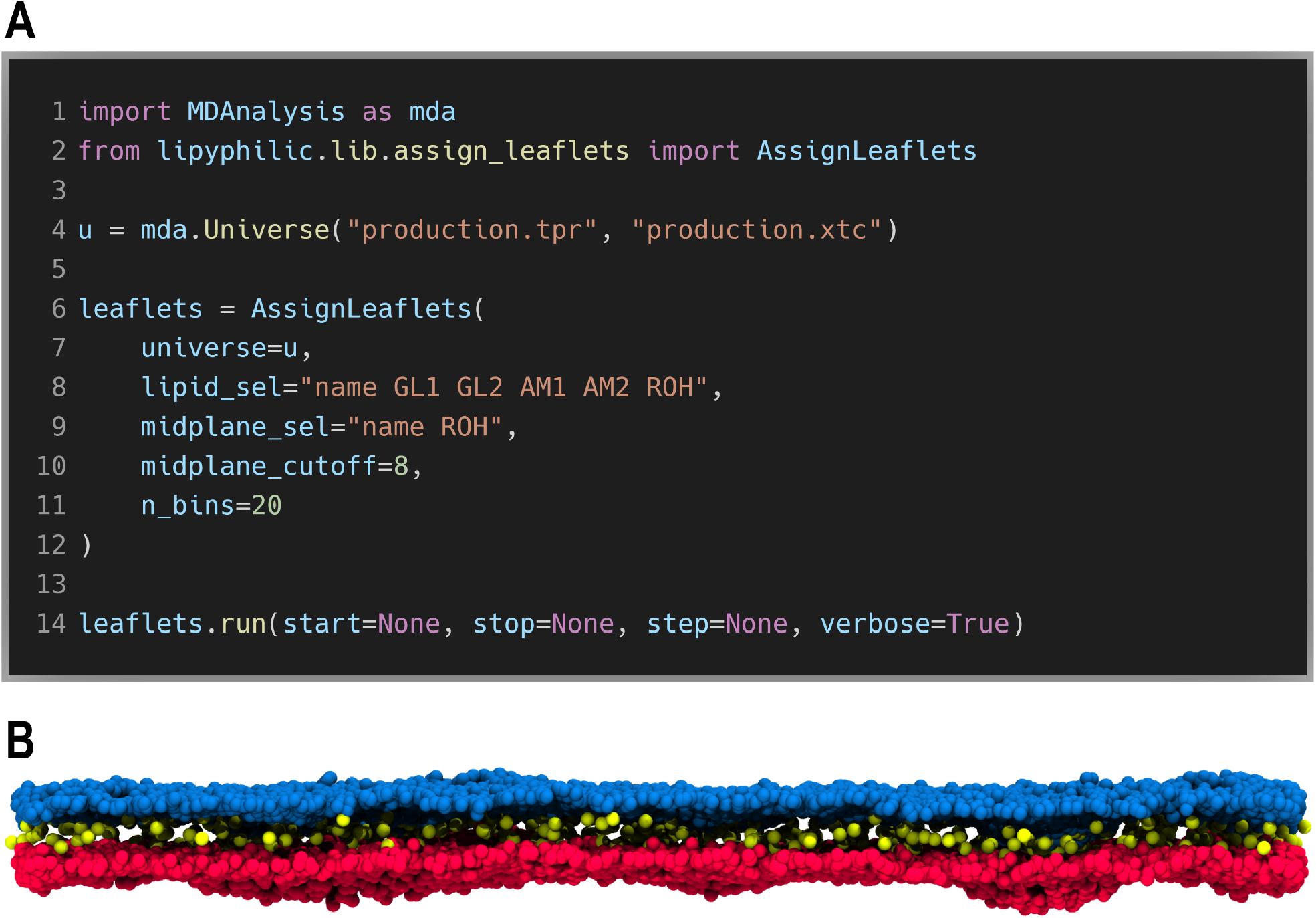
LiPyphilic can assign lipids to the upper leaflet, lower leaflet or midplane. **A**. Workflow for assigning leaflets. **B**. Lipids in the neuronal plasma membrane studied by Ingólfsson et al.^31^ are assigned to the upper leaflet (blue), lower leaflet (red) or midplane (yellow).

After creating `leaflets` as described above, the analysis is performed by calling the `run` method. Here the `start`, `stop` and `step` arguments are used to specify which frames of the trajectories to use, and a progress bar can be displayed on the screen by setting `verbose=True`. Leaflet data are then stored in the `leaflets.leaflets` attribute as a two-dimensional NumPy array. Each row in the results array corresponds to an individual lipid and each column to an individual frame. For example, `leaflets.leaflets[i, j]` contains the leaflet membership of lipid *i* at frame *j*. `leaflets.leaflets[i, j]` is equal to 1 if the lipid is in the upper leaflet, *−*1 if the lipid is in the lower leaflet, or 0 if the lipid is in the midplane.

### 2.2 Flip flop

Cholesterol is unevenly distributed across the plasma membrane, although the precise distribution is still under debate.^48^ This uneven distribution plays an important role in numerous cellular processes, and is maintained through the ultrafast spontaneous translocation, or flip-flop, of cholesterol across leaflets.

With recent advances in computing power, sterol flip-flop can now be studied directly using either coarse-grained, united-atom, or all-atom simulations. Such simulations can be used to study the flip-flop process itself or to extract rates directly from the number of observed flip-flop events. Below we describe a general analysis tool in LiPyphilic for identifying such flip-flop events in MD simulations.

The class `lipyphilic.lib.flip_flop.FlipFlop` can be used to identify successful and aborted flip-flop events. Figure 3A illustrates how to do so using the output from `AssignLeaflets`. The same MDAnalysis Universe that is used for assigning leaflets is passed to `FlipFlop`. An atom selection that specifies which molecules to consider when identifying flip-flop events is passed to the `lipid_sel` argument. The leaflet membership of each lipid selected by `lipid_sel` is passed to the `leaflets` argument. In this example, this is achieved by filtering the results array of `AssignLeaflets` to include only the leaflet membership of cholesterol molecules. The leaflet membership must be a NumPy array, of shape ((*N*_*lipids*_, *N*_*frames*_)), in which each element is equal to:

**Figure 3:**
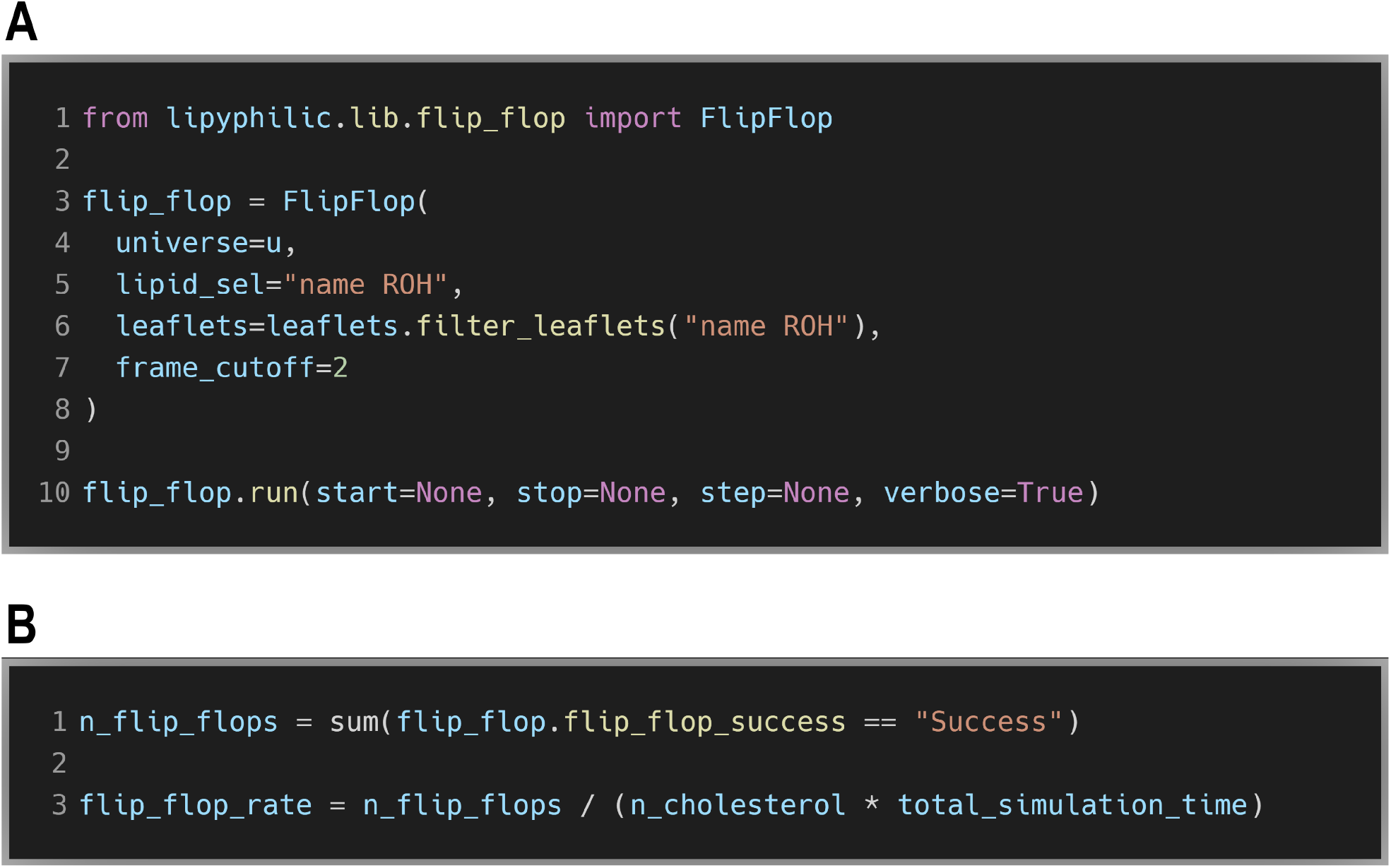
**A**. Identify all cholesterol flip-flop events, based on the leaflet membership of cholesterol at each frame. **B**. Determine the cholesterol flip-flop rate directly from the number of flip-flop events.

- 1 if the lipid is in the upper/outer leaflet
- -1 if the lipid is in the lower/inner leaflet
- 0 if the lipid is in the midplane

In this example, the `frame_cutoff` argument is used to specify that a molecule must remain in its new leaflet for at least two consecutive frames in order for the flip-flop to be considered successful.

We again call the `run` method to perform the analysis. For each molecule, LiPyphilic will then identify the frames at which it leaves one leaflet and enters another for at least `frame_cutoff` frames. If the new leaflet is different to the original leaflet, the flip-flop was successful.

The success or failure of each flip-flop event is stored in a one-dimensional NumPy array, accessible via the `flip_flop.flip_flop_success` attribute. Elements in this array are strings, equal to either “Success” or “Failure”. From this results array, it is easy to calculate the flip-flop rate based on the number of observed events, the number of cholesterol molecules in the membrane, and the total simulation time (Figure 3B).

The example in Figure 3A shows how to use the results from `AssignLeaflets` to identify flip-flop events. However, leaflets need not be identified using LiPyphilic in order to use the `FlipFlop` analysis tool. `FlipFlop` expects a NumPy array of leaflet membership as described above. Flip-flop events can thus be found even if leaflets were assigned using, for example, FATSLiM^1^ or user-written scripts^49^ based on the MDAnalysis LeafletFinder tool.

Gu et al. showed that translocation is highly influenced by the local lipid environment of a sterol.^24^ `FlipFlop` therefore not only returns the success or failure of each event, but also the frame at which the flip-flop process begins and ends along with the residue index of the flip-flopping molecule. This information is stored as a two dimensional NumPy array in the `flip_flop.flip_flops` attribute, where each row corresponds to an individual event and each column contains the:

- residue index of the flip-flopping molecule
- frame at which the molecule leaflet its original leaflet
- frame at which the molecule entered its new leaflet
- numerical identifier of its new leaflet: 1 for the upper leaflet and -1 for the lower leaflet

This information enables further analysis, such as a consideration of the local lipid environment before and after translocation.

Cholesterol is typically the only molecule to flip-flop during an MD simulation. However, ceramides and diacylglycerols, as well as fatty acids such as docosahexaenoic acid, also have fast flip-flop rates. To find flip-flop events of a molecule other than cholesterol, simply change the lipid selection passed to `FlipFlop`. For example, to find all flip-flop events for ceramides in the neuronal plasma membrane, change `lipid_sel=“resname ROH”` to `lipid_sel=“resname ?? CE”` and pass the corresponding leaflet membership NumPy array to the `leaflets` argument. This again makes use of the powerful selection language of MDAnalysis, and the fact that all ceramides in the MARTINI force field have residue names that are four characters long and end in ‘CE’.

### 2.3 Registration

The translocation of cholesterol across leaflets is thought to be important in several cellular process, including the modulation of the lateral heterogeneity of the membrane. ^47^ Recently, transient nanodomains of *L*_*O*_ phase lipids were observed in live mammalian cell plasma membranes.^50^ These nanodomains are thought to be enriched in sphingomyelin and cholesterol, and to act as functional platforms for cell signaling. However, their nature, formation, and roles in cellular processes are still not fully understood.

There is particular interest in understanding under what conditions nanodomains in apposing leaflets are spatially aligned. Such alignment is known as interleaflet registration. This has been the subject of several MD simulations, which have revealed registration to be a complex process.^11,34–42^ Registration is modulated by many factors, including the length and saturation of lipid tails as well as the relative affinity of cholesterol for the different lipid species in a domain-forming mixture.

The class `lipyphilic.lib.registration.Registration` can be used to quantify the degree of interleaflet registration in a planar bilayer. `Registration` is an implementation of the registration analysis described by Thallmair *et alii*.^40^ The degree of registration is calculated as the Pearson correlation coefficient of molecular densities in the upper and lower leaflets. First, the two-dimensional density of each leaflet is calculated:

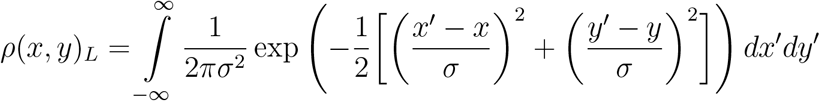

where the (*x, y*) positions of lipid atoms in leaflet *L* are binned into two-dimensional histograms with bin lengths of 1 Å. *L* is either the upper (*u*) or lower (*l*) leaflet. The two-dimensional density is then convolved with a circular Gaussian density of standard deviation *σ*. The registration between the two leaflets, *r*_*u/l*_, is then calculated as the Pearson correlation coefficient between *ρ*(*x, y*)_*u*_ and *ρ*(*x, y*)_*l*_. Values of *r*_*u/l*_ = 1 correspond to perfectly registered domains and values of *r*_*u/l*_ = *−*1 correspond to perfectly anti-registered domains.

The atoms used in calculating the interleaflet registration are specified by passing selection strings to the `upper_sel` and `lower_sel` arguments of `Registration` (Figure 4). The leaflet membership of all atoms in the two selections must be passed to the `leaflets` argument. As before, this must be a two-dimensional NumPy array of shape (*N*_*lipids*_, *N*_*frames*_). The results are stored in the `registration.registration` attribute as a one-dimensional NumPy array of length *N*_*frames*_. The array contains the Pearson correlation coefficient of the two-dimensional leaflet densities at each frame.

**Figure 4:**
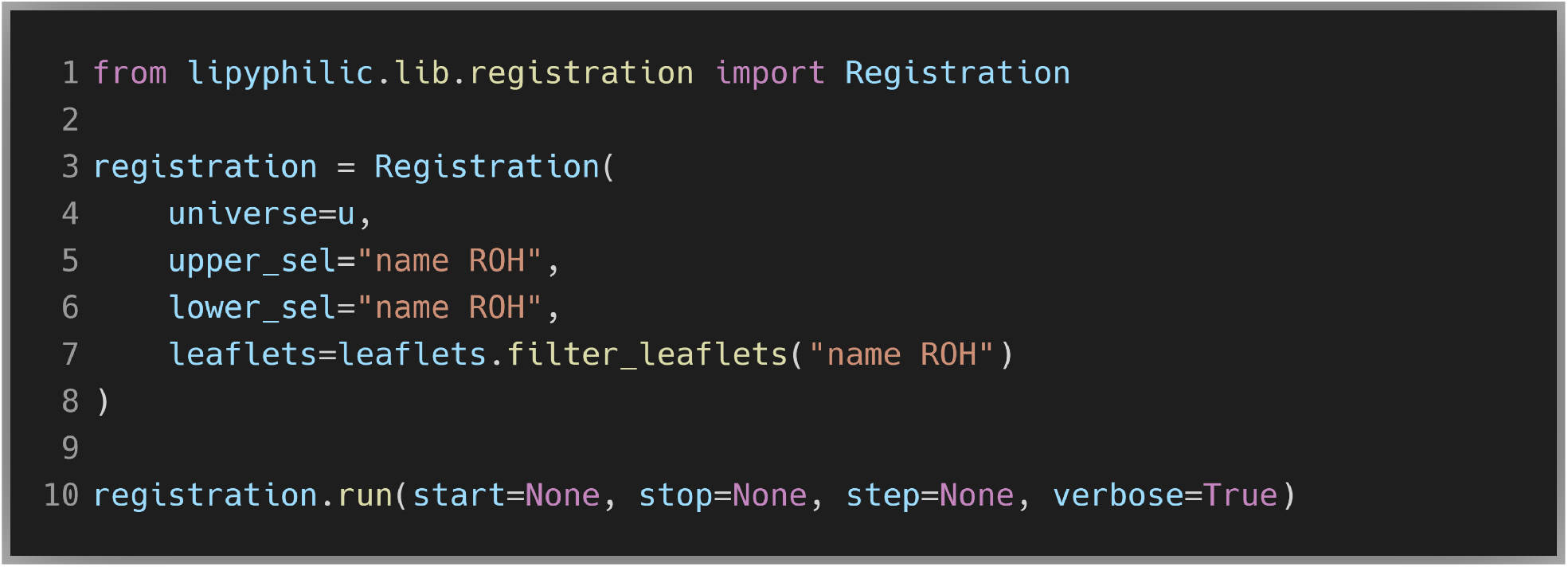
Calculate the interleaflet registration of cholesterol at each frame.

The example in Figure 4 demonstrates how to compute the registration of cholesterol across the upper and lower leaflets. However, in simulations of phase-separating mixtures it is useful to know the degree of registration of *L*_*o*_ domains, rather than the registration of a specific molecular species. `Registration` can be used to calculate this, providing the phase of each lipid at each frame has been determined using for example, ML-LPA,^5^ a hidden Markov Model,^27,33^ or the deuterium order parameter of lipid tails. If the lipid phase data is stored in a two-dimensional NumPy array of shape (*N*_*lipids*_, *N*_*frames*_), it can be used to create a boolean mask that will tell `Registration` which lipids to include in the analysis. For example, if our array is named `lipid_phase_data`, and its elements are strings of either `“Lo”` or `“Ld”`, then we can select the *L*_*o*_ lipids for analysis by passing the boolean mask `lipid_pahse_data == “Lo”` to the `filter_by` argument of `Registration`.

### 2.4 Neighbour matrix

The plasma membrane is comprised of hundreds of different lipid species. In this complex mixture, lateral heterogeneities and aggregates of specific lipid species arise spontaneously. Over the past decade, this compositional complexity has begun to feature in MD simulations of membranes.^17,25,31,51–56^ In these simulations, the lateral organization of the membrane is typically quantified via a consideration of local lipid environments. Specifically, the lipid enrichment index of species *B* around species *A, E*_*AB*_, may be defined as:^17^

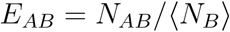

where *N*_*AB*_ is the number of molecules of species *B* around species *A*, and ⟨ *N*_*B*_ ⟩ is the mean number of species *B* around any species.

The class `liyphilic.lib.neighbours.Neighbours` provides methods for computing the lipid enrichment index and for identifying the largest cluster of a specific species of lipids over time. Both of these analyses first require the construction of an adjacency matrix, *A*, that describes whether each pair of lipid molecules are neighbouring one another or not. Two lipids are considered neighbours if they have any atoms within a given cutoff distance, *d*_*cutoff*_, of one another. The adjacency matrix can be created by passing an atom selection and a value of *d*_*cutoff*_ to the `lipid_sel` and `cutoff` arguments, respectively, of `Neighbours` (Figure 5A). The `run` method is called to construct an adjacency matrix for the frames of the trajectory specified using the `start`, `stop`, and `step` arguments. The results are available in the `neighbours.neighbours` attribute as a two-dimensional SciPy sparse matrix of shape (*N*_*Lipids*_, *N*_*lipids*_*N*_*frames*_). This matrix can be used for further analysis, either via helper methods of `Neighbours` or via user-written scripts.

**Figure 5:**
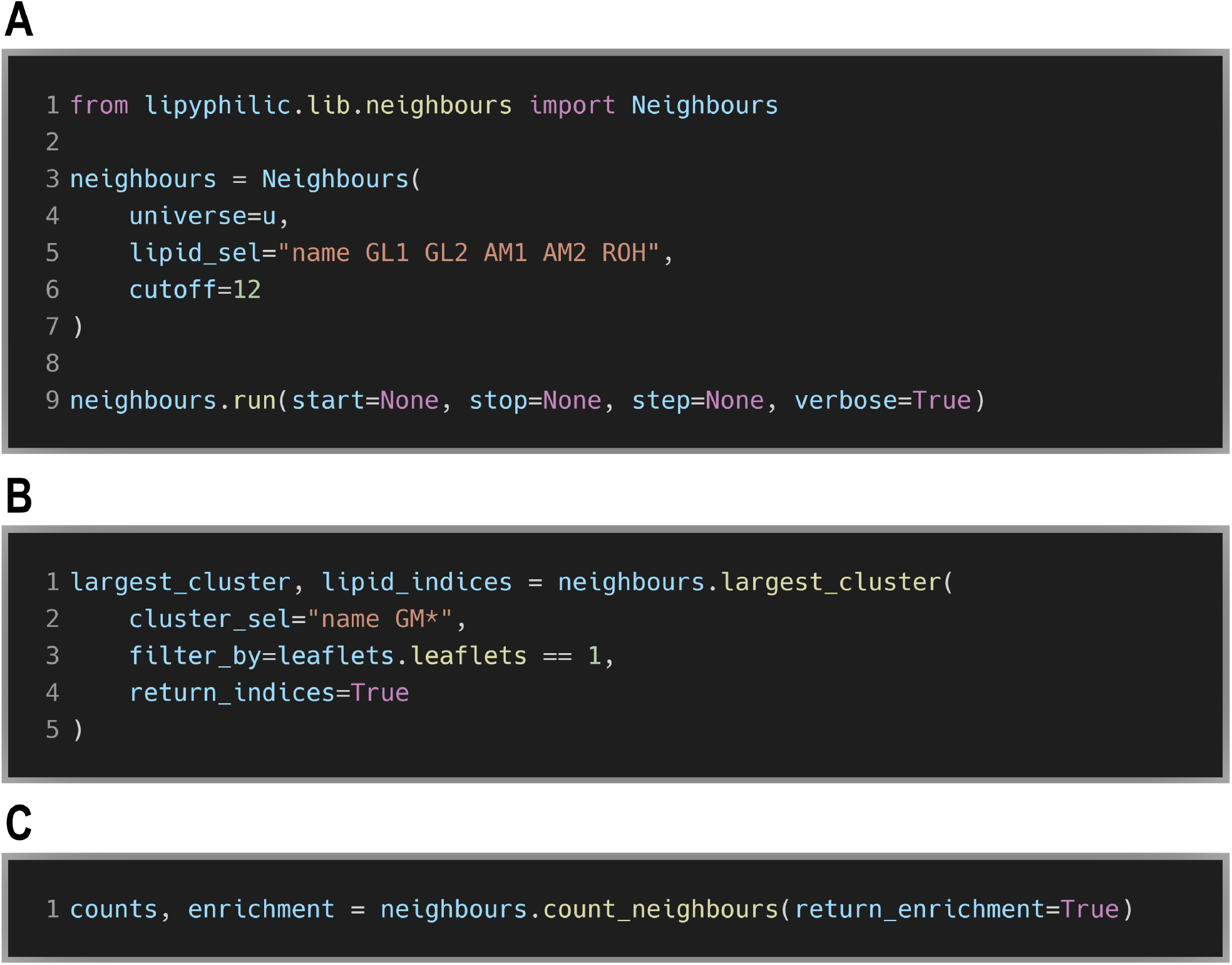
**A** Create the neighbour adjacency matrix. **B** Find the largest cluster of glycolipids at each frame as well as the residue indices of lipids in the largest cluster. **C** Calculate the enrichment/depletion index of each lipid species.

`Neighbours` initially constructs the output matrix as a SciPy `lil_matrix`, a row-based list-of-lists sparse matrix. At the end of the analysis, this is converted into a SciPy `csc_matrix`. This is because the `lil_matrix` is efficient at adding new neighbours on-the-fly, whereas the `csc_matrix` is efficient for slicing columns of the matrix. The latter is used frequently in the `largest_cluster` and `count_neighbours` helper methods of `Neighbours`.

#### 2.4.1 Largest cluster

Some glycolipids in the plasma membrane are known to aggregate, forming platforms for cell-signalling.^57–59^ The size of the largest cluster of glycolipids in the neuronal plasma membrane^31^ can be calculated using the `largest_cluster` method of `Neighbours` (Figure 5B). For this, an atom selection must be provided to the `cluster_sel` argument. In the example in Figure 5B, it is specified that only lipids in the upper leaflet should be included in the calculation by passing a boolean mask to the `filter_by` keyword. The `return_indices` argument is used to specify that the residue indices of the lipid molecules in the largest cluster at each frame are also to be returned. There is no need to call a `run` method nor to specify which frames of the analysis to use — the same frames specified in 5A will be used for the cluster analysis.

The results are not stored in an attribute in our `neighbours` object. Instead, the largest cluster size and the residue indices of the lipid molecules in the largest cluster are each returned as a NumPy array. The former is a one-dimensional array containing the number of lipids in the largest cluster at each frame. The latter is a list of NumPy arrays. Each array in the list corresponds to a single frame and contains the residue indices of the lipid molecules in the largest cluster at that frame. Knowing the indices of the lipids in the largest cluster allows for further analysis, such as calculating the lateral diffusion coefficient of lipid molecules in the cluster.^34^

To find the largest cluster at a given frame, the `neighbours.neighbours` sparse adjacency matrix is first sliced to give a matrix for the current frame only, *A*_*frame*_. The `connected_components` function of SciPy is then used to find all connected components at the current frame. NumPy’s `unique` function, with `return_counts` set to `True`, is then used to identify the largest connected component and thus the largest cluster size.

### 2.4.2 Enrichment index

After constructing the adjacency matrix, the `count_neighbours` method can be used to determine the local environment of each lipid molecule (Figure 5C). For a single lipid, its local lipid environment is defined as the number of neighbours of each species. In the example in Figure 5C, `return_enrichment=True` is set to specify that the lipid enrichment index is also to be returned.

As with `neighbours`, the results are not stored in an attribute in our `neighbours` object. The neighbour counts and the enrichment index are each returned as a Pandas `DataFrame`. The `DataFrame` of neighbour counts contains — for each lipid at each frame — the residue name, lipid residue index, frame number, number of neighbours of each species, and total number of neighbours. The lipid enrichment `DataFrame` contains the enrichment index of each lipid species at each frame. This data can easily be used to calculate the mean enrichment of each species, or it can be plotted over time to determine whether the lateral mixing of lipids has equilibrated.

#### 2.4.3 User-defined counts

By default, the `count_neighbours` method will calculate the number of neighbouring species around each individual lipid. This is done using the residue name of each lipid. However, it is also possible to use any ordinal or string data for counting lipid neighbours. For example, the enrichment index of lipids in the neuronal plasma membrane can be calculated based on their tail saturation (Figure 6). For this, first a two-dimensional NumPy array of shape (*N*_*lipids*_, *N*_*frames*_) that contains the saturation of each lipid needs to be created (Figure S1). Then this array is passed to the `count_by` argument of `count_neighbours`, and the local lipid environment and enrichment index will be determined based on the information in this array.

**Figure 6:**
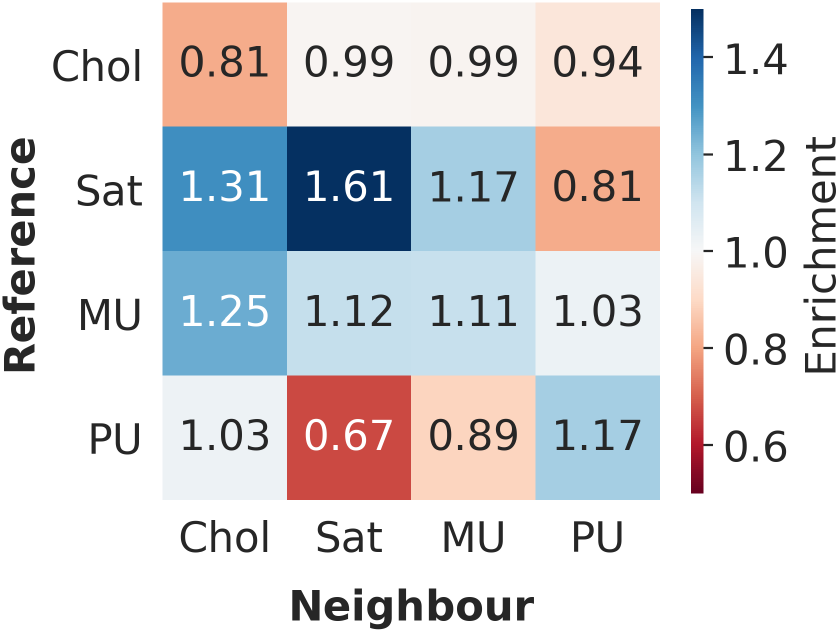
The enrichment/depletion index of lipids in the neuronal plasma membrane based on their tail saturation.

### 2.5 On-the-fly transformations

MDAnalysis has a powerful set of on-the-fly trajectory transformations. These transformations can do away with the need to create multiple instances of the same trajectory using, for example, the GROMACS `trjconv` tool. Instead, the transformations are applied each time a frame is loaded into memory by MDAnalysis. LiPyphilic extends the set of transformations available in MDAnalysis to include the ability to repair membranes split across periodic boundaries and to perform nojump trajectory unwrapping. The latter prevents an atom from being wrapped into the primary unit cell when it crosses a periodic boundary.

#### 2.5.1 Center membranes

The callable class `lipyphilic.transformations.center_membrane` can be used to center a membrane — or any supramolecular structure — in a box, providing it is not self-interacting across periodic boundaries. For each frame, all atoms in the system are iteratively shifted along a specified set of dimensions until the membrane is no longer split across the periodic boundaries (Figure 7A). After each translation, all atoms are wrapped back into the primary unit cell. For example, to check if a bilayer is split across the periodic boundary in the *z*-dimension, its extent in *z, H*, could be compared to the box length in *z, L*_*z*_. If *H* is within a user-specified cutoff value of *L*_*z*_, the bilayer is split across *z* and is thus translated in this dimension. Once the bilayer is no longer split across boundaries, it is then moved to the center of the box in *z*.

**Figure 7:**
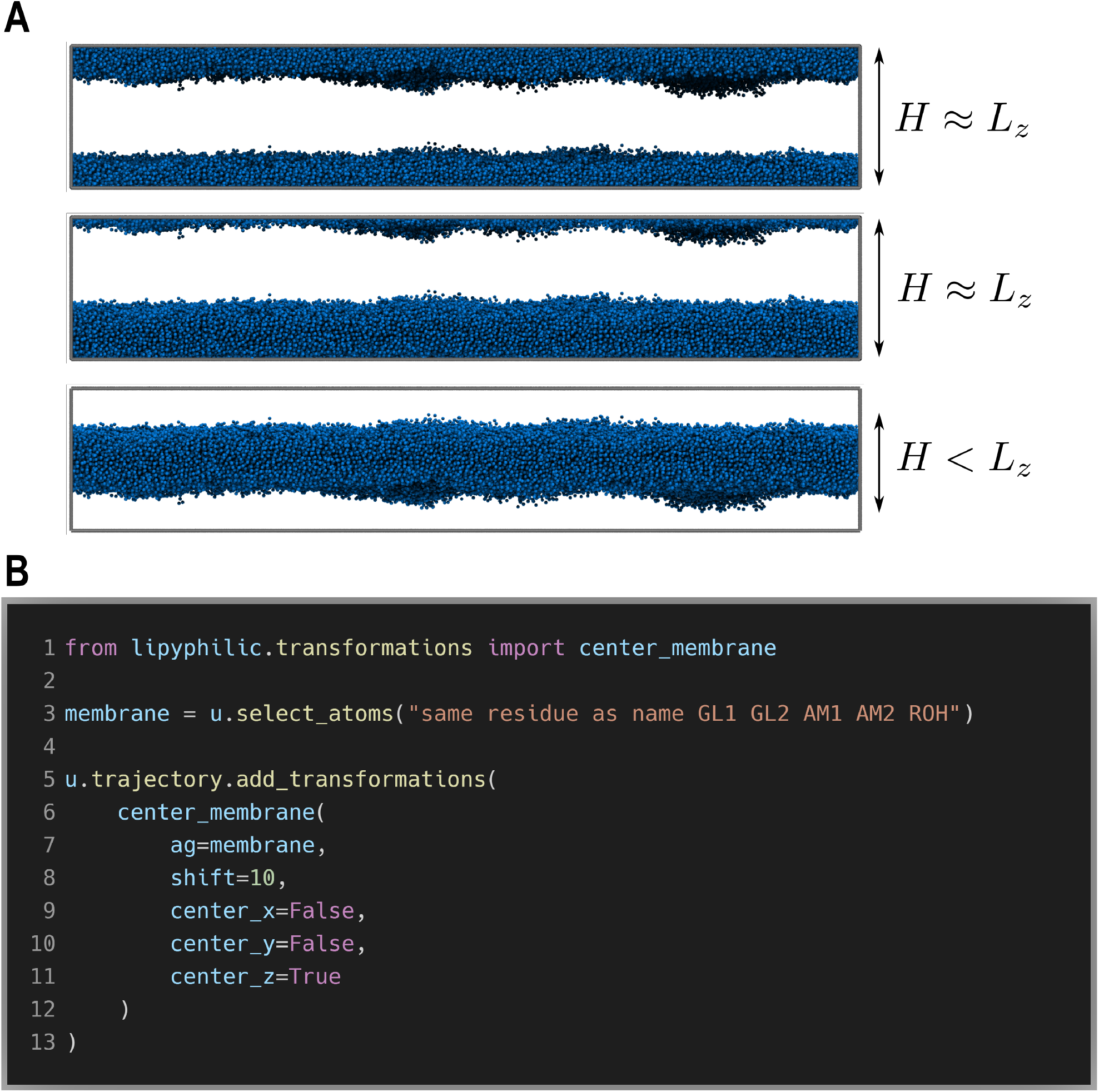
**A** A membrane split across periodic boundaries can be made whole and centered by iteratively translating the system of particles. After each translation, a check is performed to determine whether the membrane is still split across boundaries. If the extent of the membrane in *z, H*, is approximately equal to box length in *z, L*_*z*_, then the membrane is split across the periodic boundary. **B** Code snippet for applying the transformation to an MDAnalyis universe. The on-the-fly transformation can be applied to each dimension independently.

This transformation can be applied to an MDAnalysis universe, `u`, using the `u.trajectory` `.add_transformation` method (Figure 7A). This method takes as input a transformation, or a list of transformations. The `center_membrane` callable class is passed to `add_transformation` along with the arguments for `center_membrane`. The atoms that comprise the membrane are specified using the `ag` argument. The atom selection should include all atoms in the membrane, not a subset of atoms — otherwise the extent of the membrane cannot be calculated accurately. In the example in Figure 7A, the bilayer is centered in only the *z*-dimension; however, the membrane can be made whole in each dimension independently. This is controlled using the `center_x`, `center_y` and `center_z` arguments. The `shift` argument is used to specify the distance in Å that the membrane will be translated at each iteration. Too small a value of `shift` would require many iterations to make a membrane whole. Too large a value, on the other hand, may result in a membrane being translated nearly the length of the unit cell and thus remaining broken. We have found a tranlsation of 10 Å to be suitable for bilayers, but the optimal value will depend on the membrane structure and the size of the system.

#### 2.5.2 NoJump trajectory unwrapping

`lipyphilic.transformations.nojump` can be used to prevent atoms from jumping across periodic boundaries. It is analogous, but not equivalent, to using the GROMACS command `trjconv` with the flag `-pbc nojump`. This transformation can be applied to an MDAnalysis universe in much the same way as `lipyphilic.transformations.center_membrane`. We must pass an atom selection to the `ag` argument of `nojump`, and specify the dimensions to which the transformation should be applied (Figure 8).

**Figure 8:**
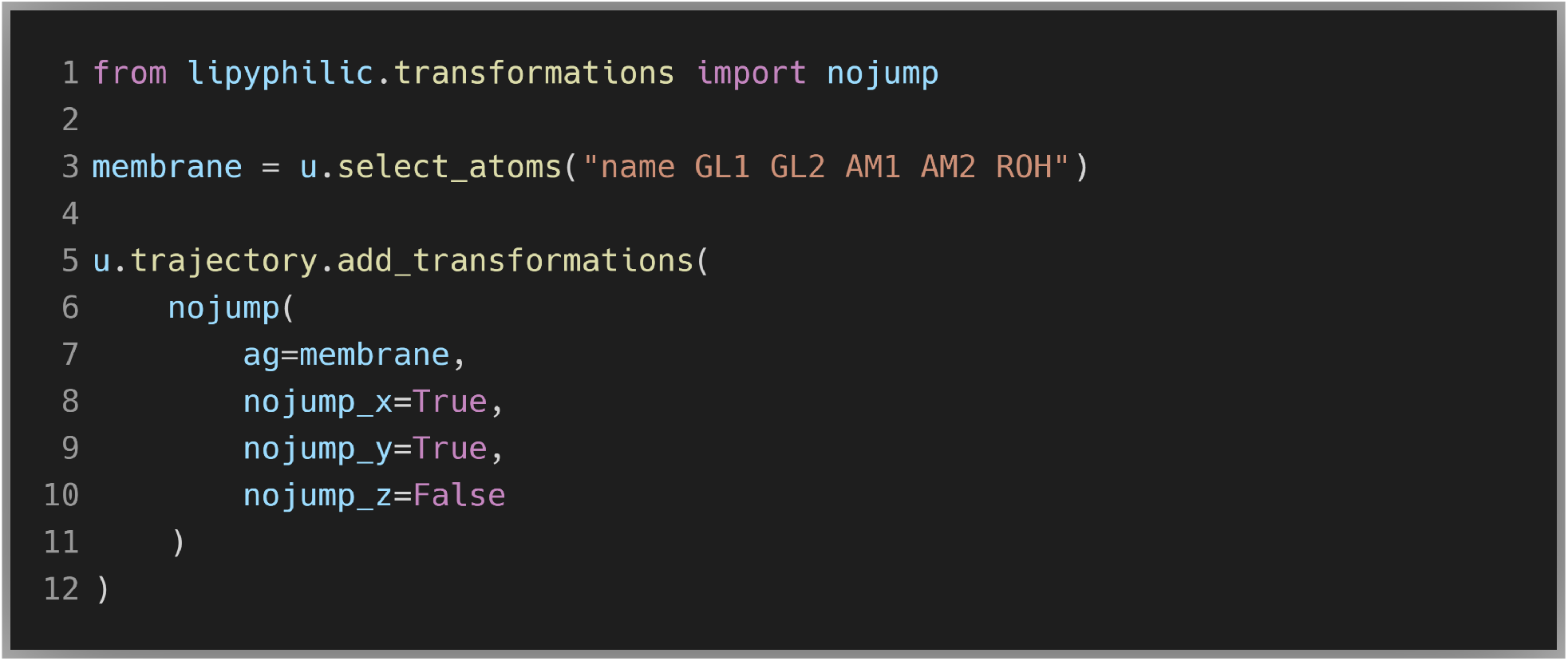
A “nojump” on-the-fly transformation can be applied to any AtomGroup in an MDAnalysis Universe. The transformation can be applied to each dimension independently.

Upon adding this transformation to your trajectory, `nojump` will perform an initial pass over the trajectory. It will determine the frames at which each atom crosses a boundary, keeping a record of the net movement of each atom at each boundary. This net movement across each boundary is used to determine the distance an atom must be translated in order to be moved from its wrapped position to its unwrapped position. Subsequently, every time a new frame is loaded into memory by MDAnalysis, such as when iterating over the trajectory, the relevant translation is applied to each atom to move it to its unwrapped coordinates.

Below we describe the nojump unwrapping algorithm implemented in LiPyphilic. We also explain how this algorithm avoids the artefacts introduced by the standard unwrapping scheme that, to our knowledge, is employed by all molecular dynamics simulation related software. Specifically, the standard unwrapping scheme fails to account for fluctuating systems sizes caused by barostats in NPT ensemble simulations.

##### Unwrapping scheme

The unwrapped position of a particle at frame *N*, denoted 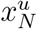, is given by:

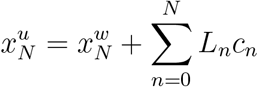

where 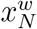 is the wrapped position of the particle at frame *N*, and 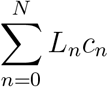 accounts for the displacement that results from all jumps across periodic boundaries from frame 0 to frame *N*. *L*_*n*_ is the box length at frame *n*, with the box centered at *L/*2. This box length is multiplied by a factor *c*_*n*_, which depends on whether an atom crossed a periodic boundary from frame *n −* 1 to frame *n*:

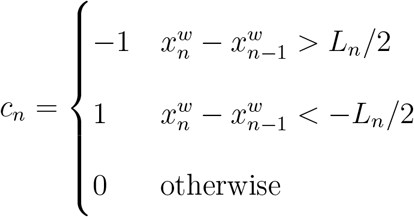

where 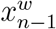 is the particle’s wrapped position at frame *n −* 1. At frame *n* = 0, for which there is no previous position, we take 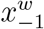 to be the particles raw atomic coordinate at frame *n* = 0. That is, if the particle is not in the primary unit cell at frame *n* = 0, we calculate the displacement required to move from 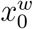 to 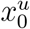.

This unwrapping scheme is equivalent to that recently described by von Bulow *et al*.,^60^ although it was derived independently. Both our unwrapping algorithm and that of von Bulow *et al*. avoid the problems that the standard unwrapping scheme suffers. To calculate 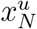, the standard unwrapping scheme iteratively adds the box length at frame *N* to the wrapped coordinated at frame *N* until 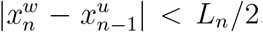. However, in the example in Figure 9, this would result in 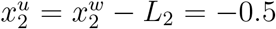 instead of the correct value 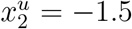. von Bulow *et al*. demonstrated clearly the effect that this inaccurate unwrapping of atomic coordinates has on the calculated diffusion coefficient.^60^

**Figure 9:**
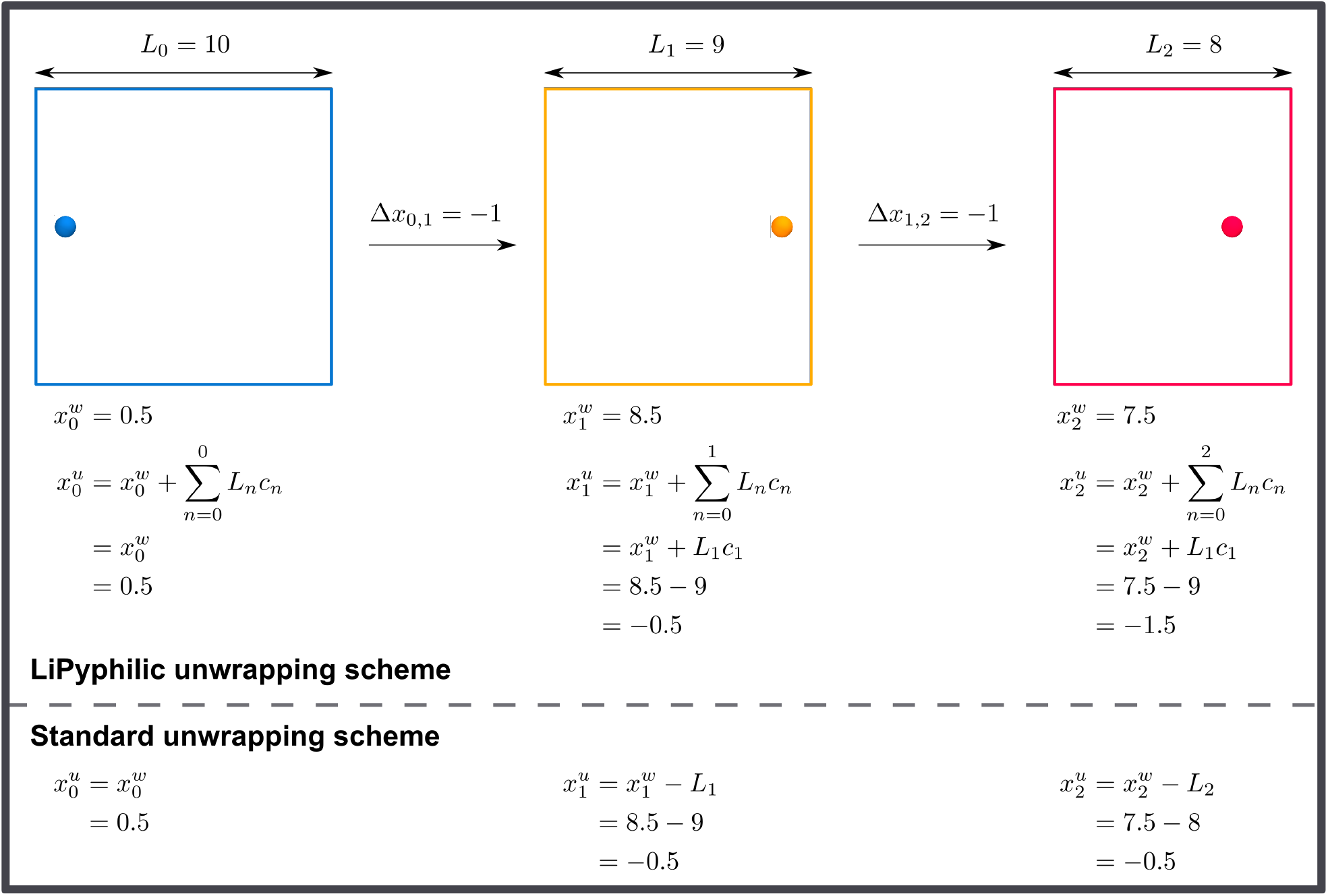
To correctly unwrap atomic coordinates we must know size of the box at the frame at which a jump across periodic boundaries occurred. The standard unwrapping scheme produces an incorrect unwrapped coordinate 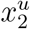.

The unwrapping scheme described above, and previously by von Bulow *et al*.,^2^ correctly accounts for the fluctuating box size in the NPT ensemble. However, it is only accurate in the case where coordinates are stored every timestep. In fact, it is impossible to correctly unwrap coordinates unless we store them at every timestep. This logically follows from the same argument made above - to correctly unwrap coordinates we must know the length of the box at the timestep at which the jump occurred. See the Supporting Information for further details.

### 2.6 Other analysis tools

While we have described some of the analysis tools that make LiPyphilic unique in the previous sections, there is much more functionality in LiPyphilic that is fully detailed in the documentation (https://lipyphilic.readthedocs.io/en/latest). This functionality includes calculating the coarse-grained lipid order parameter,^61^ the area per lipid for planar bilayers, the lateral diffusion coefficient, and the membrane thickness of planar bilayers. There are also tools for calculating thickness, orientations, and *z*-positions of lipid molecules in a bilayer. In general, LiPyphillic does not handle plotting of analysis data. However, it does have plotting utilities for visualising joint potentials of mean force (PMFs) — such as the PMF of cholesterol orientation and height — and for the projection of membrane properties onto the *xy* plane. There are full descriptions of all LiPyphilic tools, along with usage examples, in the documentation. Regarding the area per lipid tool, we recommend using either FATSLiM^1^ or MemSurfer^2^ if you have a curved membrane. These tools are designed specifically to deal with undulating bilayers and non-bilayer structures, whereas LiPyphilic will only produce reliable values in the case of planar bilayers.

## 3 Benchmarking

For benchmarking the LiPyphilic analysis tools, we have performed a simulation of an equimolar mixture of DPPC/DOPC/Cholesterol using the MARTINI 2 force field. ^62,63^ We created a symmetric bilayer of 12,000 lipids in total using the CHARMM-GUI MARTINI Maker.^64^ The production run was performed for 8.0 µs and coordinates were stored every 5.0 ns, giving a trajectory of 1,600 frames. Where analogous analysis tools are available in either FATSLiM^1^ 0.2.2 or GROMACS^65^ 2020.4, we compare their performance with that of LiPyphilic. We benchmark against FATSLiM as this is generally the fastest membrane analysis tool available.^1^ The FATSLiM benchmarks were performed using eight OpenMP threads. All other benchmarks were performed in serial. All of the benchmarks can be seen in Table 1.

**Table 1:** Benchmark times for analysis tools in LiPyphilic, using a MARTINI bilayer of 12,000 lipids. Where possible, performance is compared with either FATSLiM 0.2.2 or GROMACS 2020.4. The FATSLiM benchmarks were performed using eight OpenMP threads. All other benchmarks were performed in serial.

| Analysis/transformation | LiPyphilic class/method | Time per frame (ms) |  |
| --- | --- | --- | --- |
|  |  | LiPyphilic | Comparison |
| Leaflet identification | `AssignLeaflets` | 17.54 | 168.47<sup>†</sup> |
|  | `AssignCurvedLeaflets` | 29.85 | 168.47<sup>†</sup> |
| Flip-flop<sup>§</sup> | `FlipFlop` | 0.5 | - |
| Interleaflet registration | `Registration` | 151.81 | - |
| Construct neighbour matrix | `Neighbours` | 260.97 | - |
| Largest cluster<sup>¶</sup> | `Neighbours.largest_cluster` | 1.85 | - |
| Enrichment index | `Neighbours.count_neighbours` | 100.93 | - |
| Bilayer thickness | `MembThickness` | 16.35 | 394.11<sup>†</sup> |
| Lipid thickness | `ZThickness` | 37.62 | - |
| Lipid height | `ZPositions` | 19.05 | - |
| Lipid orientation | `ZAngles` | 22.87 | - |
| Area per lipid | `AreaPerLipid` | 1589.29 | 206.75<sup>†</sup> |
| Coarse-grained order parameter<sup>*</sup> | `SCC` | 16.10 | 119.09<sup>††</sup> |
| Unwrap membrane | `center_membrane` | 23.37 | - |
| NoJump unwarpping | `nojump` | 23.89 | 12.04<sup>††</sup> |

<sup>†</sup> FATSLiM
<sup>††</sup> GROMACS
<sup>§</sup> Time per molecule (ms) over 1,600 frames
<sup>¶</sup> Time per frame (ms) for a subset of 2,000 lipids
<sup>\*</sup> Time per frame (ms) for the sn-1 tail of DPPC (4,000 lipids)

LiPyphilic is generally very fast, with most analysis tools and trajectory transformations taking on the order of 10 ms per frame for the 12,000 lipid membrane. For example, both methods of assigning leaflets are faster than the corresponding implementation in FATSLiM. There are two further benefits of assigning leaflets with LiPyphilic: i) molecules may reside in the midplane and ii) the results are stored in a single NumPy array whereas FATSLiM creates a GROMACS index file for each frame of the trajectory. LiPyphilic is significantly faster in calculating the bilayer thickness, although the imeplmetation in FATSLiM is more sophisticated. The FATSLiM thickness tool is able to handle both bilayers and non-bilayer membranes, whereas the one in LiPyphilic is only appropriate for planar bilayers. LiPyphilic is also faster than GROMACS when it comes to calculating the coarse-grained order parameter, *S*_*CC*_. Using LiPyphilic to calculate *S*_*CC*_ is also simpler than using GROMACS, which requires the creation of a separate index group for each unique atom along a tail of each lipid species.

The calculation of interleaflet registration, construction of a neighbour matrix, and calculation of the lipid enrichment index are slower, on the order of 100 ms per frame. This is still relatively fast — although they are different analyses, these times are comparable to the performance of the various analyses available in FATSLiM such as the area per lipid and membrane thickness calculations.

The slowest analysis in LiPyphilic is the calculation of the area per lipid. This takes over 1.5 s per frame, and is 7.6 times slower than FATSLiM. We therefore recommend using FATSLiM for the area per lipid calculation if you have a large membrane (>1,000 lipids). It should be remembered, however, that the FATSLiM analyses were run in parallel over eight cores whereas all other benchmarks were performed in serial. Parallelising the analysis in LiPyphilic could result in a similar performance boost. Whilst this is out of the scope of the package in its current state, we do have plans to parallelise the slower tools in LiPyphilic in due course. In this respect, we are particularly interested in the development of Parallel MDAnalysis (PMDA).^66^ PMDA is based on MDAnalysis and uses Dask to parallelise the analysis modules.^67^ The analysis classes inherit a modified abstract base class that is specifically designed to make the parallelisation straightforward. We will wait until PMDA is out of the alpha stage of development before assessing which modules would benefit from being parallelised.

Finally, the on-the-fly transformations are fast, with `center_membrane` and `nojump` taking 23.37 ms and 23.89 ms per frame, respectively. Upon applying the `nojump` transformation, a first pass over the trajectory is performed to calculate the translations that need to be applied at each frame. This first pass takes 13.98 ms per frame for the 12,000 atoms selected in the benchmark. After this first pass, the time to load a frame into memory and apply the translations is 9.91 ms. This performance is put into perspective by considering that iterating over the trajectory with MDAnalysis takes 8.54 ms per frame itself, with no transformations applied. The total time per frame of `nojump` (23.89 ms) is approximately twice that required by the GROMACS `trjconv` tool to do the same transformation. However, using `nojump` has two benefits over GROMACS `trjconv` in that it i) prevents the need to create duplicate trajectories and ii) accounts for box size fluctuations caused by barostats.

## 4 Software Engineering

LiPyphilic is free, open-source software licensed under the GNU General Public License v2 or later. In developing LiPyphilic, we have followed the best practices in modern scientific software engineering.^68^ We use version control, unit testing and continuous integration, and we have a fully documented API with examples of how to use each analysis tool.

The full development history and planned improvements of the project are available to view on GitHub, at https://github.com/p-j-smith/lipyphilic. LiPyphilic loosely follows the GitHub-flow model of software development^69^ — developing directly from the master branch and releasing new versions soon after new functionality or fixes are added. We encourage users to submit feature requests and bug reports via GitHub, and we welcome any question about usage on our discussion page at https://github.com/p-j-smith/lipyphilic/discussions.

Unit testing in LiPyphilic is performed using Pytest.^70^ We have constructed a set of toy systems for testing each analysis tool. These systems are typically composed of two sets of atoms each arranged on a hexagonal lattice in *xy*, with the two lattices separated vertically in *z*. This setup approximates the topology of the headgroups of a lipid bilayer. Using these toy systems, we know what the results of each analysis tool should be *a priori*. We can thus test each analysis tool with full confidence in the results if the tests pass, without relying on regression tests that involve highly complex systems. Further, using Pytest-cov, ^71^ we have ensured that all analysis tools and trajectory transformations have 100% test coverage.

Finally, LiPyphilic is simple to install. We have packaged LiPyphilic to make it available for installation using widely-used package managers. The easiest way to install LiPyphilic along with all of its dependencies is through Anaconda.^72^ Alternatively, it can be installed via the Python Package Index using Pip.^73^

## 5 Conclusion

We have developed a new Python package for the analysis of lipid membrane simulations. We have focused on providing functionality not available in other membrane analysis tools, such as calculating the lipid enrichment/depletion index, the degree of interleaflet registration, and the flip-flop rate of molecules between leaflets. LiPyphilic is a modular object-oriented package that makes extensive use of NumPy,^43^ SciPy,^44^ and MDAnalysis^45,46^ for efficient computation. For analysing a 12,000 lipid MARTINI membrane, the analysis classes typically take on the order of 10-100 ms per frame. This is comparable to the performance of analysis tools in GROMACS^65^ and FATSLiM^1^ — a very fast package for membrane analysis. All analysis tools in LiPyphilic share the same API as those of MDAnalysis. This shared API makes LiPyphilic simple to learn for current users of MDAnalysis.

The modularity of LiPyphilic, along with its focus on integrating with the wider scientific Python stack, means the output of other analysis tools such as FATSLiM^1^ or ML-LPA^5^ can be used as input for further analysis in LiPyphilic. Further, the output of LiPyphilic is in the form of NumPy arrays, Scipy sparse matrices, or Pandas Dataframes. This means the results can readily be plotted or further analysed using the standard libraries of the scientific Python stack.

LiPyphiic is built upon sound software engineering principles. It uses version control, is fully unit-tested, employs continuous integration, and has extensive documentation. LiPyphilic is also trivial to install — it can be installed using either Anaconda or Pip.^72,73^ We encourage users to submit feature requests and bug reports via GitHub, and are always open to new contributors to the project.

## Acknowledgement

Via our membership of the UK’s HEC Materials Chemistry Consortium, which is funded by EPSRC (EP/L000202/1, EP/R029431/1, EP/T022213), this work used the ARCHER UK National Supercomputing Service (http://www.archer.ac.uk) and the UK Materials and Molecular Modelling Hub (MMM Hub) for computational resources, which is partially funded by EPSRC (EP/P020194/1, EP/T022213) to carry out the MD simulations of DP-PC/DOPC/Cholesterol reported in this manuscript. P.S. acknowledges the funding provided by the EPSRC DTP Studentship Block Grant (EP/N509498/1). C.D.L. acknowledges the supportive research environment of the EPSRC center for Doctoral Training in Cross-Disciplinary Approaches to Non-Equilibrium Systems (CANES, No. EP/L015854/1).

## Supporting Information Available

Python script for calculating the lipid enrichment index based on tail saturation. Discussion of problems with nojump trajectory unwrapping.

