## Supporting Information for "LiPyphilic: A Python toolkit for the analysis of lipid membrane simulations"

Figure S1: Python script for calculating the lipid enrichment index based on tail saturation.

Section S1: Problems with nojump trajectory unwrapping

```python
# Determine the saturation of each individual lipid
saturation = []
membrane = u.select_atoms("name GL1 GL2 AM1 AM2 ROH")

for lipid in np.unique(membrane.resnames):

    lipid_residues = membrane.residues.atoms.select_atoms(f"resname {lipid}").residues
    lipid_atoms = lipid_residues[0].atoms

    # It is cholesterol if it has an ROH bead
    if "ROH" in lipid_atoms.names:
        saturation.append("C")

    else:

        num_doulbe = sum([True for name in lipid_atoms.names if name.startswith("D")])

        # It is saturated if it has no D beads
        if num_doulbe == 0:
            saturation.append("S")

        # It is monounsaturated if it has 1 D bead
        elif num_doulbe == 1:
            saturation.append("M")

        # It is polyunsaturated if it has 2+ D beads
        else:
            saturation.append("P")

saturation = np.asarray(saturation)

# Now create a two-dimensional NumPy array where each lipid is labelled by its saturation
count_by = np.full((membrane.n_residues, neighbours.n_frames), fill_value="", dtype=str)
for lipid, sat in zip(np.unique(membrane.resnames), saturation):

    lipid_indices = membrane.residues.resnames == lipid
    count_by[lipid_indices, :] = sat

# Finally, count the negihbours based on tail saturation
counts, enrichment = neighbours.count_neighbours(
    count_by=count_by,
    return_enrichment=True
)
```

Figure S1: Workflow for calculating the enrichment index of lipids in the neuronal plasma membrane[1] based on their degree of tail saturation. This assumes the neighbour adjacency matrix has already been constructed as shown in Figure 5A.

### More trajectory unwrapping problems

The unwrapping scheme described in the main text, and previously by von Bulow *et al.*,[2] correctly accounts for the fluctuating box size in the NPT ensemble. However, it is only accurate in the case where coordinates are stored every timestep. In fact, it is impossible to correctly unwrap coordinates unless we store them at every timestep. This logically follows from the same argument made in the main text - to correctly unwrap coordinates we must know the length of the box at the timestep at which the jump occurred.

This can be seen when considering Figure 9, and assuming that coordinates are written every other frame. In this case, the coordinates are only known at timesteps $N = \{0, 2\}$ and so:

$$x_N^u = x_N^w + \sum_{n=0}^{N/2} L_{2n} c_{2n}$$

where $x_N^w$ is the wrapped position of the particle at frame $N$, and $\sum_{n=0}^{N/2} L_{2n} c_{2n}$ accounts for the displacement that results from all jumps across periodic boundaries from frame 0 to frame $N$, determined using coordinates stored *every other* frame. $L_{2n}$ is the box length at frame $2n$ and:

$$c_{2n} = \begin{cases} -1 & x_{2n}^w - x_{2n-2}^w > L_{2n}/2 \\ 1 & x_{2n}^w - x_{2n-2}^w < -L_{2n}/2 \\ 0 & \text{otherwise} \end{cases}$$

In this example, using the atomic coordinates in Figure 9:

$$
\begin{aligned}
x_2^u &= x_2^w - \sum_{n=0}^{N/2} L_{2n} c_{2n} \\
&= 7.5 - 8.0 \\
&= -0.5
\end{aligned}
$$

which would give a displacement from frame 0 to frame 2 of −1, rather than −2. Therefore, the length of the box must be known at the frame at which the jump actually occurred. If this information is not known, then artifacts may be introduced into the unwrapping of atomic coordinates, even using the unwrapping scheme described here and by von Bulow *et al.*[2]
